# The molecular genetics of hand preference revisited

**DOI:** 10.1101/447177

**Authors:** Carolien G.F. de Kovel, Clyde Francks

## Abstract

Hand preference is a prominent behavioural trait linked to human brain asymmetry. A handful of genetic variants have been reported to associate with hand preference or quantitative measures related to it. Most of these reports were on the basis of limited sample sizes, by current standards for genetic analysis of complex traits. Here we performed a genome-wide association analysis of hand preference in the large, population-based UK Biobank cohort (N=331,037). We used gene-set enrichment analysis to investigate whether genes involved in visceral asymmetry are particularly relevant to hand preference, following one previous report. We found no evidence implicating any specific candidate variants previously reported. We also found no evidence that genes involved in visceral laterality play a role in hand preference. It remains possible that some of the previously reported genes or pathways are relevant to hand preference as assessed in other ways, or else are relevant within specific disorder populations. However, some or all of the earlier findings are likely to be false positives, and none of them appear relevant to hand preference as defined categorically in the general population. Within the UK Biobank itself, a significant association implicates the gene *MAP2* in handedness.

## Introduction

Hand preference is a conspicuous behavioural trait, with about 10% of people preferring to use their left hand for many tasks, about 1% having no preference, and the large majority preferring to use the right hand ^1^. Hand preference is probably initiated during prenatal phases, and further established in early infancy ^2-5^. Gene expression analysis has revealed left-right differences in the human central nervous system as early as four weeks post conception ^6,7^, which indicate that laterality is an innate and pervasive property of the central nervous system. Across different regions of the world and throughout history, the behavioural variation has persisted, albeit that cultural suppression of left-handedness has sometimes occurred ^8-11^.

There has been much interest in possible causes of hand preference, as well as its associations with cognitive variation and disorders ^12-15^. The existence of a genetic component for hand preference has been investigated in large twin studies, consisting of thousands of individuals. These studies found additive heritabilities of around 25% for left-hand preference ^16-19^. In terms of individual genetic loci, monogenic models for hand preference were originally conceived, such as the dextral/chance model ^20^. The model proposed that there are two alleles, D (dextral) and C (chance), with the homozygous DD genotype producing right-handers (directional asymmetry), the homozygous CC genotype producing random asymmetry (50% right-handers and 50% left-handers) due to a hypothesized loss of direction-setting in brain development, and the heterozygote, DC, being intermediate and producing 25% left-handers and 75% right-handers. Another related idea, conceived with respect to continuous measures of left-versus-right hand motor skill, was the ‘right-shift’ model, which proposed a major gene whose alleles cause either rightward mean asymmetry, or no directional bias ^21^.

However, genetic linkage analyses in families with elevated numbers of left-handed members have not found convincing evidence for single major genes ^22-24^. Genome-wide association scan (GWAS) analysis has also not found any genomic locus with a large enough effect on hand preference to be compatible with monogenic models, unless there would be extensive genetic heterogeneity (i.e. many different Mendelian effects involved in the population)^25^. Meanwhile, as analyses based on twins have indicated 25% heritability, there is clearly ample room for environmental influences, although these remain mostly unknown. Early life variables such as birthweight and twinning can influence handedness, but not to an extent which is remotely predictive at the individual level ^26,27^.

Various molecular genetic studies have identified individual genes as potential contributing factors to handedness, with small effects, which would be consistent with a polygenic architecture underlying the heritable component to this trait. We review these genes briefly here:

- A sib-pair analysis (N=89 sibships with reading disability) found suggestive linkage of left versus right relative hand skill, a continuous measure based on the pegboard task ^28^, to a broad region on chromosomes 2 ^29^. A follow-up study suggested the signal came from paternal inheritance ^30^. Further analysis suggested *LRRTM1* (leucine rich repeat transmembrane neuronal 1) as the most likely gene, as it showed evidence for paternal-specific association with relative hand skill of a particular haplotype, as well as imprinted expression in cell lines ^31^.
- A CAG-repeat polymorphism in the gene AR (androgen receptor), located on chromosome X, has been studied in relation to hand preference, due to a longstanding theory that brain laterality is influenced by sex hormones ^32^. One study found association effects which were opposite in males and females for the CAG-repeat ^33^. A different study found that right-handedness was associated with shorter CAG repeats in both sexes ^34^, while another, which only included males, found that mixed-handedness was associated with longer CAG repeats ^35^.
- In a set of families affected by bipolar disorder, an association was found of a variant in *COMT* (catechol-O-methyltransferase) with relative hand skill, although this did not survive multiple testing correction ^36^. *COMT* was studied because it was considered a candidate gene for bipolar disorder ^37^.
- A study of the genes *SETDB1* (SET domain bifurcated 1) and *SETDB2* (SET domain bifurcated 2) found that the single nucleotide polymorphism (SNP) rs4942830 in *SETDB2* was significantly associated with hand preference as measured as a quantitative trait on the Edinburgh scale ^38^ (N=950 healthy people), although for the binary trait of left versus right-handedness, the p-value did not survive multiple testing correction ^39^. This result was replicated, including the stronger effect for the quantitative trait (right=732, left=75, healthy people)^40^. *SETDB2* was chosen as a candidate for study as it has been shown to regulate structural left-right asymmetry in the zebrafish central nervous system ^41^.
- A GWAS study of relative hand skill, in a sample of people with reading problems (3-stage design, N=728), found a genome-wide significant association in the gene *PCSK6* (proprotein convertase subtilisin/kexin type 6) ^42^. The same research group performed a second GWAS in a cohort of individuals from the general population (N=2666), and found no genome-wide significant associations, and no indication of involvement of *PCSK6* ^43^. The GWAS results also yielded evidence that genes influencing visceral asymmetry might be involved in relative hand skill in both the dyslexic group and the general population group. For this analysis, the authors tested sets of genes for an enrichment of association signals in the GWAS data, where the sets were defined based on mutations in mice that influence the organization of the visceral organs on the left-right axis ^43^.
- In a GWAS study of 263 left vs 2092 right-handers, and 173 left vs 1412 right-handers (this was a study of healthy twins, and the GWAS analysis was divided to achieve non-relatedness of subjects), no genome-wide significant results for self-reported hand preference were found ^25^. The most significant associations were near the neighbouring genes *SCN11A/TTC21A/WDR48* (sodium voltage-gated channel alpha subunit 11/tetratricopeptide repeat domain 21A/ WD repeat domain 48), and *AK3*/*RCL1* (adenylate kinase 3/ RNA terminal phosphate cyclase like 1) with p<1E-6, i.e. a suggestive level of association which does not survive statistical correction for multiple testing over the whole genome.
- A GWAS of 4268 individuals from the general population, using a questionnaire-based definition of handedness, found no association p-values below 5E-06 in the genome ^44^. Possibly other negative results remained unpublished.

It has become increasingly clear during the last decade that psychiatric, cognitive and behavioural traits are usually influenced by many genetic factors with very small effects of each individual locus, so that tens of thousands of subjects are needed to detect effects reliably ^45^. None of the studies listed above approached that level of sample size or statistical power. Low power reduces the likelihood that a statistically significant result reflects a true effect ^46^. Most of the studies summarized above made secondary use of datasets which were collected for different purposes, so that left-handedness was present at only roughly 10% within the datasets. This meant that statistical power was even less than might have been achieved in such sample sizes, for example through selection schemes balanced for handedness. Given the statistical issues affecting the earlier studies, it seems likely, from a contemporary perspective on statistical genetics, that most or all of the findings were false positives.

Some of the genetic associations summarized above were identified in groups with specific disorders, including children with reading disability ^29,42^, and families with bipolar disorder ^36^. Therefore, even if the genetic associations were real, they may not extrapolate to the general population. Finally, a variety of different measures related to handedness were used in these studies, including binary traits based on simple questions such as ‘which hand do you write with’, to quantitative indices based on the peg board test.

An opportunity arose recently to perform a high-powered GWAS of hand preference in a very large, adult population sample from the UK, known as the UK Biobank dataset ^47^. This dataset comprises more than 300,000 subjects with handedness information, assessed by questionnaire (see below). SNP-based heritability analysis, based on variation over all of the autosomes (i.e. all chromosomes minus the sex chromosomes) arrived at an explained variance (SNP-heritability) for left-handedness of only 1.8% in the UK Biobank (p=6.4e-03), which is markedly lower than previous indications from twin studies (see above), while ambidexterity had an estimated SNP-heritability of 6.8% (p=8.7e-04)^48^.

Here we performed GWAS for hand preference in this large and well powered dataset, to identify novel loci associated with this trait, and to investigate whether any of the previously reported loci, summarized above, show evidence for association with binary trait handedness in the general adult population. We also applied different methods of gene-set enrichment analysis to the GWAS results, in order to test the relevance of gene-sets that are involved in laterality of the visceral organs.

## Methods

### Data

We used the data collected in the UK Biobank ^47^ and provided to Clyde Francks as part of research application 16066. Signed informed consent was obtained from all participants, and the UK Biobank has ethical approval from the UK Biobank Research Ethics Committee under number 16/NW/0274. All methods were performed in accordance with the relevant guidelines and regulations. For descriptions of all traits in the UK Biobank cohort, see http://biobank.ctsu.ox.ac.uk/crystal/search.cgi.

The phenotype 1707 ‘Handedness’’ was based on a touchscreen question “Are you right or left handed?”, with four answering options: ‘right-handed’, ‘left-handed’, ‘use both right and left hands equally’, and ‘prefer not to say’. We merged ‘right-handed’ and ‘use both right and left hands equally’ into the category ‘non-left-handed’ to create a dichotomous phenotype. ‘Prefer not to say’ was treated as missing data.

In a previous study we identified various early life factors which affect handedness in the UK Biobank dataset^49^. For the purposes of the present study, some of these were relevant to use as covariate effects when performing genetic association analysis: sex (trait 31), year of birth (trait 34), country of birth (trait 1647), and being part of a multiple birth (trait 1777). The traits ‘birthweight’ (trait 200022) and ‘being breastfed’ (trait 1677) also showed association with hand preference^49^, but we did not include them here as covariates, as they have significant SNP-heritabilities of their own in the UK Biobank dataset (https://nealelab.github.io/UKBB_ldsc/). In this case, some genetic loci may conceivably influence both handedness and one of these traits, while we wished to detect any genetic influences on hand preference in our GWAS without correcting for shared genetic effects. In addition, as covariates we used the first 20 principal components which describe genome-wide diversity within the dataset, as provided by the UK Biobank^50^, in order to control for population stratification. We restricted the analysis to participants with British ancestry, as reported before^50^. Individuals from this restricted dataset were selected for the present study when they had data for the handedness phenotype and covariates used. Within this subset, we excluded randomly one individual from each pair of participants whose genetic relatedness was inferred to be 3rd degree or closer, on the basis of genome-wide genotype data, as previously calculated ^50^, individuals were removed when there was a mismatch of their reported and genetically inferred sex, putative aneuploidy of the sex chromosomes, excessively high genome-wide heterozygosity (> 0.19) or genotype missingness (missing rate >0.05). The total number of participants remaining was N= 331,037: left-handed= 31,856, non-left-handed = 299,181, the latter comprised of right-handed = 293,857 and ambidextrous = 5,324. The GWAS for left-handed versus non-left-handed was our primary analysis, the results of which we used for downstream analyses at the whole gene level and gene-set level (below) However, in addition, we ran GWAS for the trait right-handed vs non-right-handed and ambidextrous vs non-ambidextrous, for look-ups of previously reported individual SNPs (see Introduction).

Imputed genotype data ^50^ were available as dosage data for ∼97 million SNPs per individual. QCtool (version: 2.0.1, revision 872b463, http://www.well.ox.ac.uk/∼gav/qctool_v2/index.html) was used to assess SNP quality, and SNPs were retained if in the selected subsample the imputation ‘info’ score was > 0.7, the minor allele frequency (MAF) was > 0.001, and the P value for the Hardy-Weinberg equilibrium (HWE) test was > 1E-07. This left ∼15 million SNPs. GWAS analysis was run using linear regression assuming an additive genetic model, with BGENIE v1.2 ^50^.

When we ran GWAS for ‘ambidextrous’ versus ‘non-ambidextrous’, due to the much smaller size of the ambidextrous group than the other handedness groups, we raised the MAF threshold to 0.01.

### Previously implicated SNPs and genes

We checked the GWAS catalog (http://www.ebi.ac.uk/gwas/search, 3 July 2018) for the terms ‘handedness’ and ‘hand’ to find SNPs and genes previously implicated in traits related to hand preference. In addition, we checked the literature through Pubmed (July 2018) for ((‘hand preference’ OR ‘hand skill’ OR ‘handedness’) AND genetic) to identify SNPs or genes found in non-GWAS studies (such as candidate gene studies). We also checked references in the papers we found. The results of this search are summarized in the Introduction.

In the results of our GWAS for hand preference in the UK Biobank, we first checked the p-values for association, for each of the specific SNPs reported in the previous papers (Table 1). We also generated association scores for each individual gene implicated in the previous papers, by use of the Magma program as implemented in the FUMA package v1.3.3c (see Supp. Table 1 for specific settings) ^51^(November 2018). Each gene-based score summarizes the overall level of association at all SNPs spanning a given gene, into one single score and generates a corresponding P value for that entire gene.

**Table 1.** Significance of association with hand preference measures in the UK Biobank, for SNPs and genes previously implicated in the literature.

| GENE | CHR | START | END | SNP | P-GENE<sup>a</sup> | P-SNP<br>LEFT<sup>a</sup> | P-SNP<br>RIGHT<sup>b</sup> | P-SNP<br>AMBI<sup>c</sup> | REF |
| --- | --- | --- | --- | --- | --- | --- | --- | --- | --- |
| **LRRTM1** | 2 | 80515483 | 80531874 | rs1007371 | 0.42 | 0.81 | 0.74 | 0.80 | <sup>31</sup> |
| **PCSK6** | 15 | 101840818 | 102065405 | rs7182874 | 0.22 | 0.97 | 0.90 | 0.69 | <sup>43</sup> |
| **COMT** | 22 | 19929130 | 19957498 | rs4680 | 0.78 | 0.95 | 0.89 | 0.64 | <sup>36</sup> |
| **SETDB2** | 13 | 50018429 | 50069138 | rs4942830 | 0.52 | 0.45 | 0.94 | 0.11 | <sup>39</sup> |
| **SCN11A** | 3 | 38887260 | 38992052 | rs883565 | 0.80 | 0.14 | 0.09 | 0.47 | <sup>25</sup> |
| **AK3** | 9 | 4711155 | 4742043 | rs296859 | 0.77 | 0.98 | 0.92 | 0.76 | <sup>25</sup> |
| **AR** | X | 66764465 | 66950461 | micro-<br>satellites<sup>d</sup> | 0.76 | - | - | - | <sup>33</sup> |

**Notes:** Gene-level association is shown for the left-handed versus non-left-handed phenotype, and SNP level association for all three hand preference contrasts. START and END are according to the reference genome build 37 (see methods).<sup>a</sup> Using GWAS of left-handed vs non-left-handed. <sup>b</sup> Using GWAS of right-handed vs non-right-handed. <sup>c</sup> Using GWAS of ambidextrous vs non-ambidextrous. <sup>d</sup> The original study made use of a microsatellite repeat polymorphism which was not genotyped or imputed in the UK Biobank dataset.

### Gene-sets involved in visceral laterality

To test for the involvement of genes associated with visceral asymmetry in relative hand skill, Brandler et al.^43^ had selected a number of mouse phenotypes related to visceral asymmetry (see Introduction), such as ‘dextrocardia’ and ‘situs inversus’, from all those available in the Mouse Genome Informatics (MGI) resource (http://www.informatics.jax.org/). In this database, mutations in specific genes have been linked to resulting phenotypes, such that each phenotype has a set of individual genes annotated to it. The exact criteria by which Brandler et al. selected their specific phenotypes for analysis were not described. In addition, new relevant mouse laterality phenotypes, and their associated genes, may have been added since 2013. We therefore made an up-to-date selection of phenotypes related to visceral asymmetry in the MGI, in the following way: In the hierarchical tree for mammalian phenotypes (http://www.informatics.jax.org/vocab/mp_ontology, MGI 6.12, last updated 06/19/2018), we searched for ‘heterotaxy’, ‘situs inversus’, ‘isomerism’ and ‘dextrocardia’, and also included all child terms if they also related to asymmetry phenotypes. We only included terms with at least 15 genes. This approach resulted in a list of 11 mouse phenotypes (See Results). The mouse genes were then mapped to corresponding human genes, using homologene.data (v68, May 2014) from ftp://ftp.ncbi.nih.gov/pub/HomoloGene/current, by matching mouse genes with human genes that had the same HID (HomoloGene group identifier). The list of phenotypes and the associated human genes can be found in Supplementary Table 2.

In the Online Mendelian Inheritance in Man database OMIM (https://omim.org/, last updated on 26 Jun 2018), we also identified all genes in the human phenotypic series ‘Primary Ciliary Dyskinesia’ (MIM #244400), because half of the patients with this disorder present with situs inversus. These genes were used to form one additional gene-set, thus in this study we tested a total of twelve sets, eleven from the mouse database and one based directly on human genetics (See Results).

Seven of our mouse phenotypes were among those previously analysed by Brandler et al. (Supp. Table 3). We also repeated our analyses using the exact same list of mouse phenotypes used by Brandler et al. (Supp. Table 3). However, the gene lists associated with each mouse phenotype may have changed since Brandler et al. performed their study five years ago, as new information on genes affecting visceral asymmetry can have been incorporated.

## Gene-set enrichment analysis

We ran three different gene-set enrichment algorithms that have been developed for application to GWAS results. Brandler et al., in their investigation of gene-sets involved in visceral laterality, had used MAGENTA ^52^. For consistency we also used MAGENTA (v2, 21 July 2011), plus two newer algorithms called Magma (^53^, v1.06) and Pascal (^54^, no version number). The three enrichment algorithms work in different ways, but essentially they first assign a single association score to each gene based on the GWAS data, and then test whether the genes in a given set show, on average, more evidence for association with the trait in question than the rest of the genes in the genome for which scores could be calculated (hence these are sometimes called ‘competitive’ analyses ^55^), while accounting for non-independence of SNPs due to linkage disequilibrium. For Magma and Pascal, we used default settings, while for MAGENTA we chose the settings as used by Brandler *et al.* (parameter settings for all analyses can be found in Supp. Table 1). Pascal and Magma offered the option to calculate gene-scores based on the single most associated SNP within a gene (the ‘max’ or ‘top’ option), or on a combination of SNPs within the gene (the ‘sum’ option), and we ran both approaches with both softwares, for a robust assessment of whether the previously implicated gene-sets in the literature might affect handedness in the UK Biobank.

## Results

None of the SNPs previously reported in the literature, in datasets other than the UK Biobank itself (see Introduction), had a significant effect on left-handedness in the UK Biobank (Table 1). Nor was there any effect at the whole-gene-level, for any of the previously reported genes (Table 1). Also for the trait definitions of ‘ambidextrous vs non-ambidextrous and ‘right-handed vs non-right-handed’, none of the previously claimed SNPs from other datasets showed significant association in the UK Biobank (Table 1).

The gene-set enrichment analyses did not convincingly support the proposition that sets related to visceral asymmetry are particularly associated with left-handedness (Table 2, Supp. Table 3). There was one potential association involving the gene-set ‘right-sided isomerism’ in the analysis with MAGENTA, but this was not supported by any of the other methods.

**Table 2.** Gene-set enrichment results for visceral asymmetry-related gene-sets, based on the UK Biobank GWAS for left-handedness versus non-left-handedness.

| mouse ID | Description | Pascal<br>sum P | Pascal<br>max P | Magma<br>sum P | Magma<br>max P | Magenta<br>P |
| --- | --- | --- | --- | --- | --- | --- |
| MP:0000508 | right-sided isomerism | 0.65 | 0.83 | 0.98 | 0.46 | 0.006<sup>a</sup> |
| MP:0000531 | right pulmonary isomerism | 0.64 | 0.68 | 0.97 | 0.76 | 0.06 |
| MP:0000542 | left-sided isomerism | 0.77 | 0.59 | 0.52 | 0.73 | 0.10 |
| MP:0000644 | Dextrocardia | 0.68 | 0.87 | 0.85 | 0.84 | 0.23 |
| MP:0001706 | abnormal left-right axis patterning | 0.93 | 0.84 | 0.67 | 0.93 | 0.41 |
| MP:0002766 | situs inversus | 0.40 | 0.51 | 0.70 | 1.00 | 0.49 |
| MP:0003178 | left pulmonary isomerism | 0.70 | 0.52 | 0.55 | 0.90 | 0.13 |
| MP:0004133 | Heterotaxia | 0.80 | 0.89 | 0.16 | 0.99 | 0.35 |
| MP:0010808 | right-sided stomach | 0.34 | 0.72 | 0.86 | 0.95 | 0.17 |
| MP:0011250 | abdominal situs ambiguus | 0.08 | 0.62 | 0.70 | 0.94 | 0.31 |
| MP:0011252 | situs inversus totalis | 0.23 | 0.67 | 0.71 | 0.99 | 0.44 |
| PCD | human primary ciliary dyskinesia | 0.33 | 0.31 | 0.84 | 0.70 | 0.30 |

**Notes:** Different softwares and settings were used. See Supplementary Table 1 for the settings. The uncorrected P values for each gene-set and enrichment analysis are shown. <sup>a</sup> FDR=0.035.

In the GWAS for left-handedness vs non-left-handedness in the UK Biobank dataset (Figure 1), the following significant associations were found with P values below the threshold 5E-08 for correcting for multiple testing over the whole genome: on chromosome 2q34 the SNP rs142367408 (p=3.8E-08), located 28 kilobases upstream of the gene *MAP2* (Microtubule associated protein 2); on chromosome 17q21 the SNP rs144216645 (p=1.2E-09) within a large inversion haploblock spanning the genes *STH*, *KANSL1*, *ARL17B*, *ARL17A*, *LRRC37A*, *LRRC37A2* and *NSF*; on chromosome 13q22 the SNP rs11454570 (p=4.7E-08) within an intron of the non-coding RNA-gene LINC00381.

**Figure 1.**
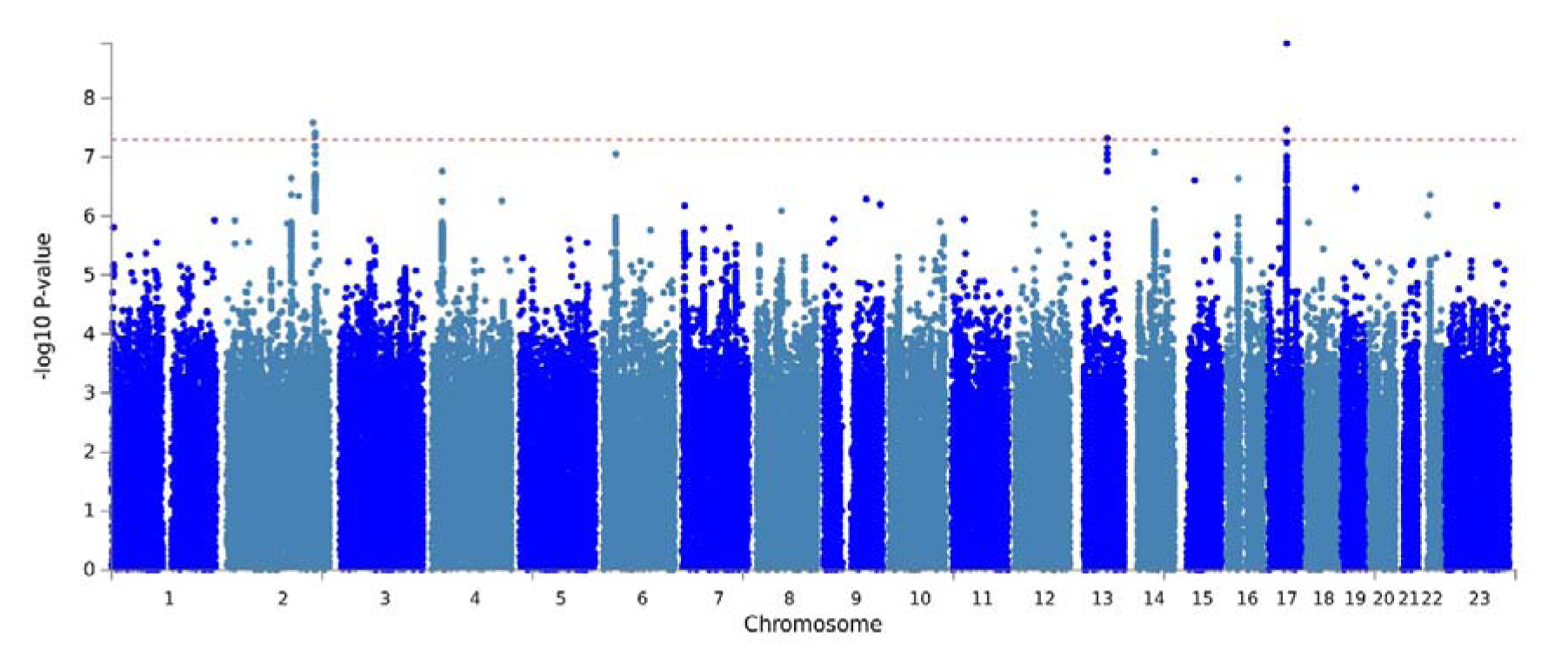
Manhattan plot of GWAS results for left-handed vs non-left-handed people in the UK Biobank sample. The red dashed line marks the genome-wide significance threshold of p=5.0E-08. (Plot produced with FUMA^51^).

The results for right-handed vs non-right-handed people were similar to those for left-handed versus non-left-handed (Supp. Figure S1), which was expected as only the relatively small group of ambidextrous people had moved between groups. For right-handed versus non-right-handed, the region around *MAP2* on chromosome 2 did not reach genome-wide significance. Instead, the major histocompatibility complex region on chromosome 6p21 showed significant associations. The most associated SNP was rs9366770 (p=2.2E-09), located in an exon of the non-coding RNA *HCG27*. The inversion region on chromosome 17 was also associated with the right-handed versus non-right-handed phenotype, again most significantly with the SNP rs144216645 (p=2.3E-08). An additional association on chromosome Xq25 was detected with the SNP rs767669906 (p=1.0E-08), located 44 kilobases downstream of the gene *DCAF12L1*. However, with a minor allele frequency of 0.0035, this signal may not be reliable.

The GWAS for ‘ambidextrous’ vs ‘non-ambidextrous’ showed no significant associations.

Zoomed in figures showing genetic association within the significantly associated regions can be found in the supplement (Figures S2-S5)^56^.

## Discussion

In a very large dataset comprising hundreds of thousands of adult subjects from the general population, we found no evidence for an involvement in hand preference of any genetic variants, genes, or gene-sets that have been previously claimed in the literature. We consider the most likely explanation to be that the earlier findings were false positives, as the study sample sizes were orders of magnitude smaller than the present analysis (see Introduction). However, as the earlier studies used various different trait measures related to handedness, and some made use of selected disorder populations, then it remains possible that some of the earlier findings were indeed real, but not detectable in the present analysis (see Introduction).

There was no evidence for an involvement, in hand preference, of common variants in genes related to visceral left-right patterning. Note that Brandler et al. ^43^ performed analysis of a quantitative handedness index based on peg moving, and included a sample with reading disability, when they originally reported such a relationship. Our present analysis was not an attempt to directly replicate the study of Brandler *et al*., but rather it was a related investigation, of genes involved in visceral asymmetry with respect to questionnaire-defined, binary-trait hand preference, in a large dataset from the general population. In the rare genetic disorder Primary Ciliary Dyskinesia (PCD)(MIM # 244400), recessive mutations in specific genes involved in ciliary biology result in a 50% chance of mirror-reversed left-right visceral asymmetry, a condition known as *situs inversus totalis* (SIT) ^57^. When PCD and SIT co-occur (the combination is known as Kartagener syndrome), it appears that the rate of left-handedness is unchanged from the general population ^58^. However, there may be an increase of left-handedness in people with SIT when they do not have PCD ^59,60^, which would suggest the existence of some developmental mechanisms which link visceral asymmetry with brain asymmetry. Nonetheless, a recent genome-wide mutation screening study did not identify any individual genes as likely candidates to cause non-PCD SIT ^61^, and it is possible that the aetiology is multifactorial or even non-genetic ^61,62^. At present, the study by Brandler et al. remains the only one to report tentative evidence for a molecular genetic link between visceral laterality and handedness.

We found that two loci, on 2q34 and 17q21, were associated with left-hand preference in the UK Biobank, as well as a third locus on chromosome 13 which had borderline significance. The most likely causative gene within the large chromosome 17q21 inversion cannot be determined on the basis of these data, as the region spans at least twelve genes showing similar levels of association with the trait (Supp. Figure S2). This region shows association with numerous other human traits (see e.g. http://big.stats.ox.ac.uk/variant/17-43659975-T-C). The variant on chromosome 13 resides in an intron of the non-coding RNA-gene LINC00381 (Supp Figure S3). This SNP is part of a short A-repeat, but is in LD with other SNPs (www.ensembl.org). For the locus on chromosome 2q34, the most probable gene appears to be *MAP2*: the association signal extends from upstream into this gene’s coding region, and the single most associated SNP is associated with expression differences of *MAP2* in oesophageal tissue, according to the GTeX database (https://www.gtexportal.org/home/ Release V7 (dbGaP Accession phs000424.v7.p2)), and not with any other gene’s expression level (Supp. Figure S4). MAP2 protein is a well-known marker for neuronal cells, and specifically the perikarya and dendrites of these cells ^63^. MAP2 is involved in neurogenesis ^64^, and almost exclusively expressed in the brain ^65^. The protein plays a role during the development of various brain structures (e.g. ^66,67^), and in mice seems to play a role in motor skills ^68^. One report suggested that *MAP2* is more highly expressed in right hippocampus than left during development in the rat ^69^. How exactly this gene may act in affecting motor laterality in human development remains unknown. Regardless, as this genetic association is based on hundreds of thousands of study participants, *MAP2* must be considered the single most reliably implicated gene in left-handedness yet identified, although replication will be desirable.

When pooling ambidextrous people together with left-handers, to create a ‘right-handed’ versus ‘non-right-handed’ phenotype, an additional locus within the MHC region of chromosome 6p was detected. Again this is a very broad region of linkage disequilibrium spanning many genes. We found that the most associated SNP is an eQTL for the non-coding RNA *HCG27* (HLA complex group 27) in the cerebellar hemisphere and frontal cortex, according to the GTeX portal. Other associated SNPs within the region, rs3130976 (p=2.0E-08) and rs2854008 (p=2.4E-08), are eQTLs for MICB (MHC class I polypeptide-related sequence B) and *C4A* (complement C4A) in the cerebellar hemisphere, but for other genes in other tissues. We cannot conclude which may be the most likely causative gene or genes in the region, on the basis of this information.

There may have been some particular sources of noise in the UK Biobank definitions of hand preference. An earlier analysis of the UK biobank dataset ^26^ suggested a reduction in left-hand preference of roughly 20% among the oldest participants (born around 1940), relative to younger participants (born around 1970), which suggests that some of the older left-handed people were forced to switch to right-handedness during childhood. Our GWAS analysis did adjust for year of birth as a covariate effect, but hand switching may have reduced power. In addition, the location of birth within the UK was found to influence the probability of being left-handed ^26^, likely primarily for cultural rather than biological reasons. Country of birth was therefore included as a covariate in our GWAS. Furthermore, the ambidextrous group was previously found to give a high rate of inconsistent answers between different assessment visits ^26^. However, ambidextrous people comprised only a relatively small subset.

It is striking that common genetic variants in the UK Biobank dataset have little overall effect on hand preference, as measured by a very low SNP-based heritability (only 2-3%)^48,70^, while previous twin-based studies have indicated a heritability of roughly 25% for left-hand preference (see Introduction). It has been noted before that behavioural traits can show a particularly large difference between twin and SNP-based heritability estimates ^71^. A commonly given explanation is that much of the heritability may be due to rare SNPs and mutations, which are not well captured by genotyping arrays and imputation protocols in GWAS studies. Rare mutations can be relatively recent in origin, and under negative selection. Whether these possibilities apply to hand preference will need to be investigated in future large-scale genome sequencing studies.

## Conclusion

In conclusion, previously reported genetic associations with measures related to handedness, found in datasets other than the UK biobank, were not supported by analysis of this very large cohort. We also found no supportive evidence that common variants within genes involved in visceral left-right patterning contribute to hand preference. We consider the most likely explanation for the discrepancy to be that the previous findings were noise, which arose from limited sample sizes. Differences of trait measurement and population selection might also be involved. Regardless, what is known of the molecular genetic basis of hand preference remains tentative and extremely limited, with the gene *MAP2* providing the most robust lead so far.

## Supporting information

## Acknowledgements

This research has been conducted, using the UK Biobank Resource. We thank all participants of the UK Biobank for their contribution. CGFdeK was supported by an Open Programme grant (824.14.005) to CF from the Netherlands Organization for Scientific Research (NWO). C.F. was supported by the Max Planck Society (Germany).

## Author contributions

CF initiated the study and edited the manuscript. CGFdK carried out and interpreted the data analysis, and wrote the manuscript.

## Additional information

The authors declare to have no conflicting interests.

### Data availability

Data were obtained from the UK Biobank cohort, as part of research application 16066, with Clyde Francks as the principal applicant. Summary and description of these data can be found at http://biobank.ctsu.ox.ac.uk/crystal/. For the use of the data, application must be made to http://www.ukbiobank.ac.uk/register-apply/.

## Supplemental Information

Supp. Table 1: Software packages and settings for the various gene-set enrichment analyses.

Supp. Table 2 (an Excel file): Gene-sets used in the current study in GMT-format. Genes have Entrez identifiers.

Supp. Table 3: Results for visceral asymmetry-related gene-sets as used in the previous analysis by Brandler *et al*. (2013)

Supp. Figures S1-S5. Further detailed plots of genetic association results.

