## Supplementary material for "The molecular genetics of hand preference revisited"

### Supplementary information

- Supplemental Table 1
- (Supplemental Table 2 is a separate Excel file)
- Supplemental Table 3
- Supplemental figures S1-S5

### Supplemental tables

**Supp. Table 1** Software and settings for the various gene-set enrichment analyses (GSEA) reported in Table 2 and Supp. Table 3.

| package/option | package version | up-stream (kb)^a^ | down-stream (kb)^a^ | MAF-cutoff^b^ | Gene-scoring^c^ | GSEA-scoring^c^ | include HLA^d^ |
| --- | --- | --- | --- | --- | --- | --- | --- |
| MAGMA IN FUMA | 1.06 | 20 | 20 | 0.01 | sum | competitive | No |
| MAGENTA^e^ | vs2_July2011 | 20 | 10 | 0 | max | competitive | No |
| PASCAL, MAX | 2015 | 40 | 15 | 0.05 | max | competitive | No |
| PASCAL, sum | 2015 | 40 | 15 | 0.05 | sum | competitive | No |
| MAGMA, max | 1.06 | 10 | 5 | 0.01 | max | competitive | Yes |
| MAGMA, SUM | 1.06 | 10 | 5 | 0.01 | sum | competitive | Yes |

^a^ The extent of inclusion of flanking regions upstream and downstream of gene definitions.

^b^ Cut-off for minor allele frequency of SNPs.

^c^ See Methods for sum vs max gene-scoring, and for competitive GSEA-scoring.

^d^ Whether the Human Leukocyte Antigen region on chromosome 6p was included (a very large region of extended linkage disequilibrium).

^e^ We report the 75^th^ percentile cutoff as explained in reference [52].

**Supp. Table 3**. Gene set enrichment results for the visceral asymmetry-related gene-sets as analyzed originally by Brandler et al. 2013 [43].

| **Mouse Phenotype ID** | **Phenotype** | **Pascal sum P** | **Pascal max P** | **Magma sum P** | **Magma max P** | **Magenta P** |
| --- | --- | --- | --- | --- | --- | --- |
| **MP:0000276** | heart right ventricle hypertrophy | 0.74 | 0.89 | 0.21 | 0.53 | 0.78 |
| **MP:0000284** | double outlet right ventricle (*) | 0.68 | 0.62 | 0.28 | 0.69 | 0.26 |
| **MP:0000508** | right-sided isomerism | 0.65 | 0.86 | 0.98 | 0.46 | 0.004^a^ |
| **MP:0000531** | right pulmonary isomerism | 0.64 | 0.72 | 0.97 | 0.76 | 0.06 |
| **MP:0000542** | left-sided isomerism | 0.77 | 0.60 | 0.52 | 0.73 | 0.10 |
| **MP:0000644** | dextrocardia | 0.68 | 0.87 | 0.85 | 0.84 | 0.24 |
| **MP:0001706** | abnormal left-right axis patterning | 0.93 | 0.84 | 0.67 | 0.93 | 0.41 |
| **MP:0002625** | heart left ventricle hypertrophy | 0.87 | 1.00 | 0.69 | 0.48 | 0.55 |
| **MP:0002766** | situs inversus (*) | 0.40 | 0.51 | 0.70 | 1.00 | 0.49 |
| **MP:0003922** | abnormal heart right atrium morphology | 0.26 | 0.16 | 0.10 | 0.59 | 0.22 |
| **MP:0004133** | heterotaxia (*) | 0.80 | 0.87 | 0.16 | 0.99 | 0.35 |
| **MP:0004158** | right aortic arch | 0.87 | 0.79 | 0.9 | 0.91 | 0.26 |
| **MP:0009569** | abnormal left lung morphology | 0.36 | 0.42 | 0.98 | 0.78 | 0.11 |
| **MP:0009570** | abnormal right lung morphology(*) | 0.36 | 0.42 | 0.98 | 0.78 | 0.11 |
| **MP:0010429** | abnormal heart left ventricle outflow tract morphology | 0.42 | 0.54 | 0.22 | 0.25 | 0.33 |

Gene set enrichment results for the same visceral asymmetry-related gene-sets as analyzed originally by Brandler et al. 2013 [43], based on the UK Biobank GWAS for left-handedness versus non-left-handedness, and using different software and settings. See Supplementary Table 1 for the settings. The uncorrected P values for each gene set and enrichment analysis are shown.

(*) adjusted p< 0.05 for the reading-disabled cohort in Brandler *et al.* (2013).

^a^ FDR-adjusted p-value 0.034.

### Supplemental figures

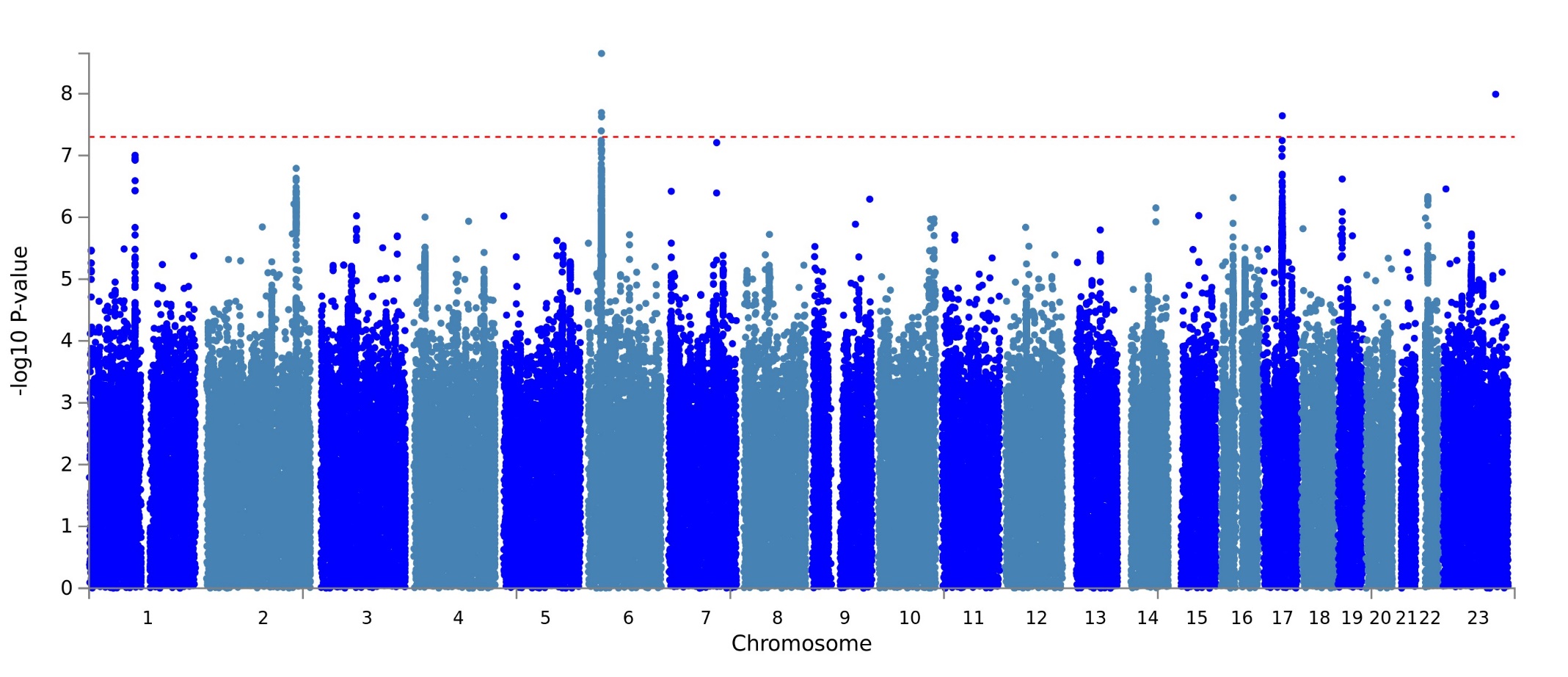

**Figure S1**. Manhattan plot of GWAS results for **right-handed** vs non-right-handed people in the UK Biobank sample. Red dashed line marks the genome-wide significance threshold of p=5.0E-08. (Plot produced with FUMA [51]).

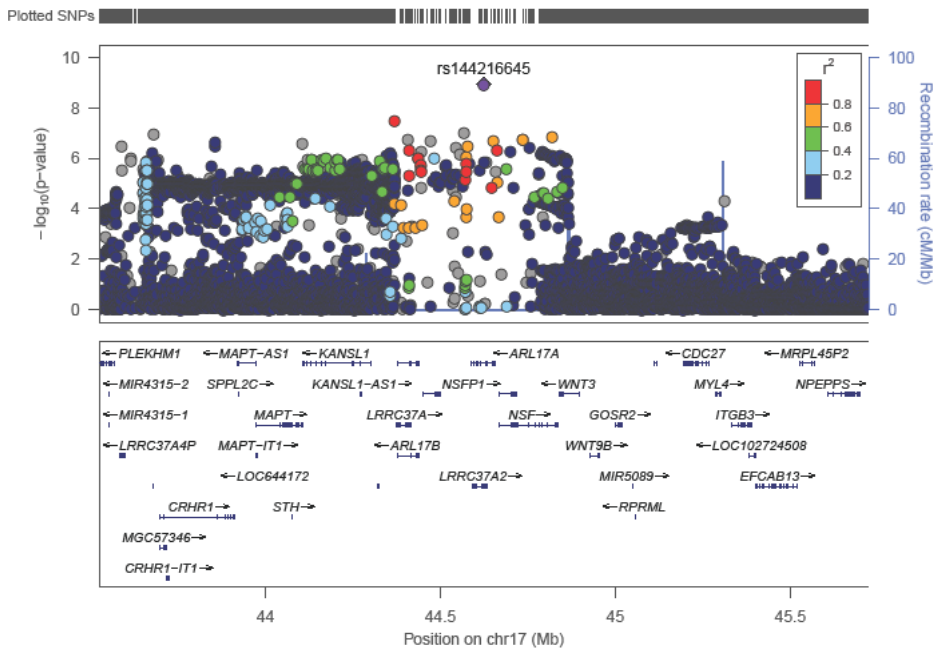

**Figure S2**. Detailed association plot for left-handed vs non-left-handed people in the associated region on chromosome 17. SNPs are coloured for their linkage disequilibrium (LD) with top hit rs144216645. (Plot produced with <http://locuszoom.org/>, Pruim J.R. et al. 2010, Bioinformatics 2010 September 15; 26(18): 2336.2337 [56]). Please note that the plots S2-S5 have X-axes of different lengths.

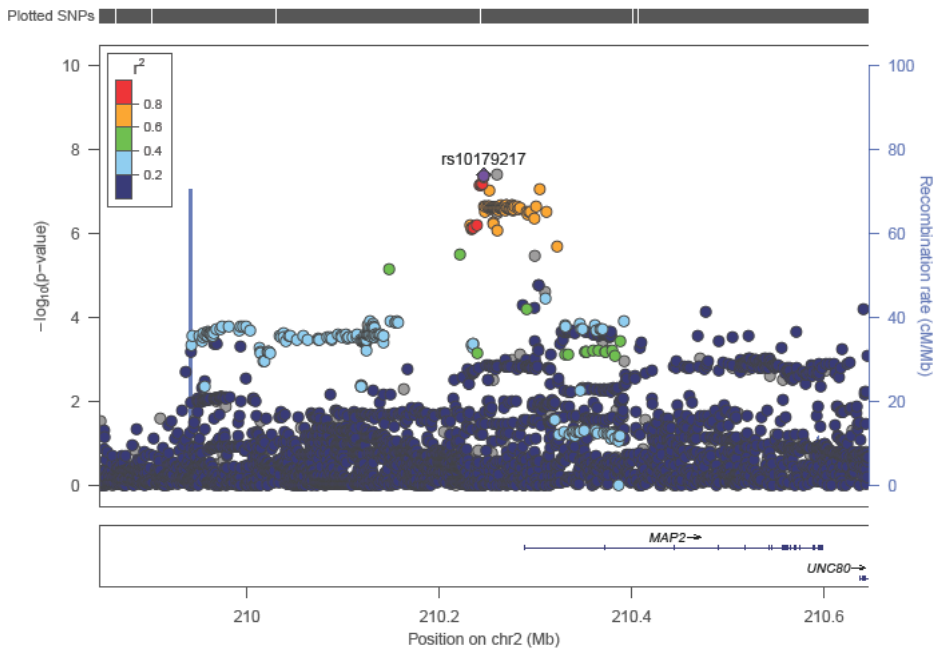

**Figure S3**. Detailed association plot for left-handed vs non-left-handed people in the associated region on chromosome 2. Because no LD data were available in Locuszoom for the top SNP rs142367408, SNPs in the plot are coloured for LD with the second most associated SNP rs10179217 (p=4.0E-08), indicated by the diamond.

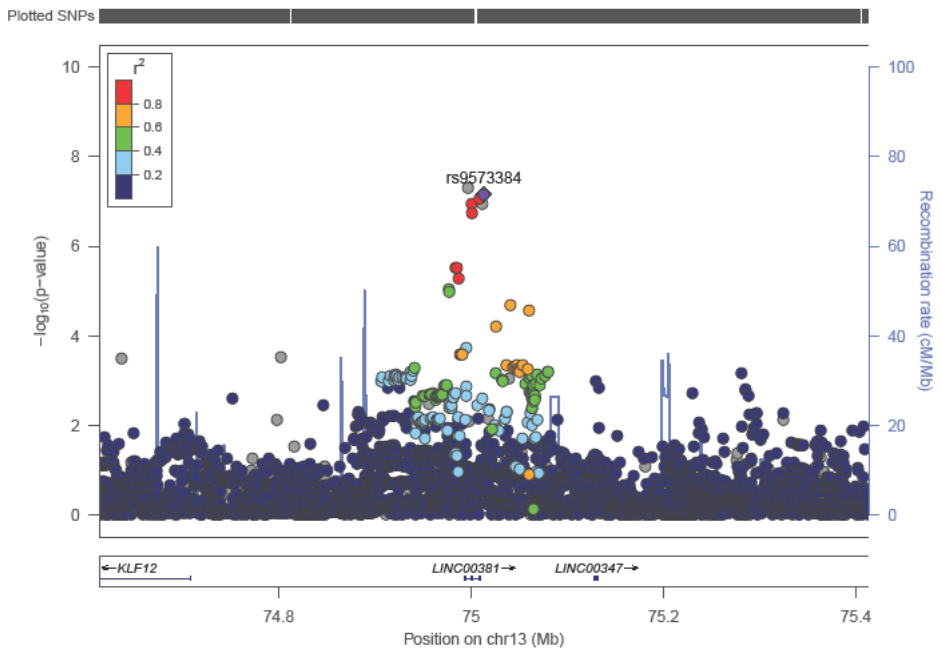

**Figure S4**. Detailed association plot for left-handed vs non-left-handed people in the associated region on chromosome 13. Because no LD data were available for the top SNP rs11454570, SNPs are coloured for LD with the second most associated SNP rs9573384 (p=6.8E-08), indicated by the diamond.

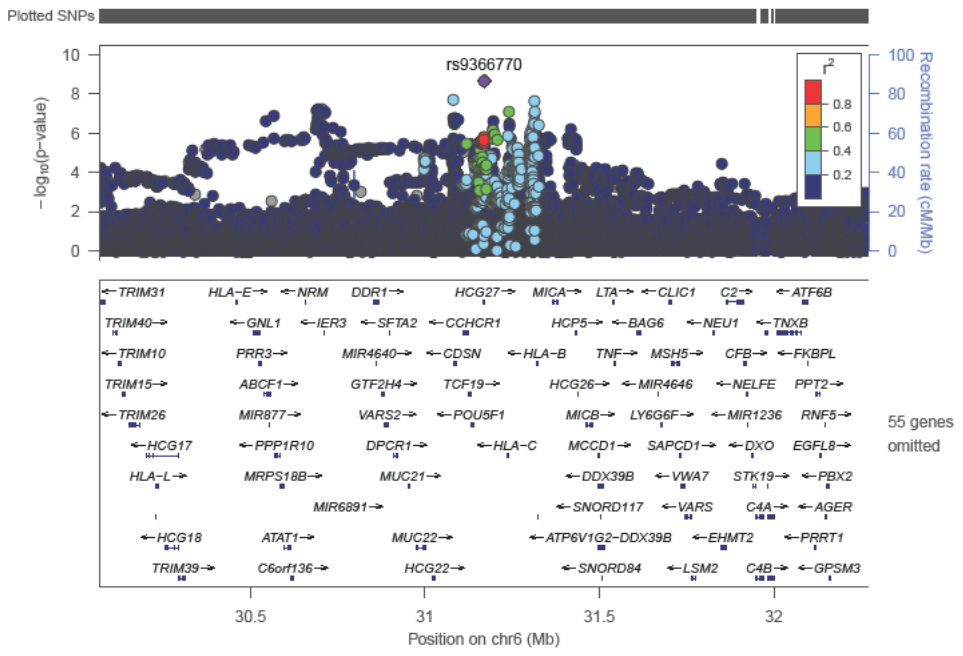

**Figure S5**. Detailed association plot for **right-handed** vs non-right-handed people in the associated region on chromosome 6, MHC region.
